# Retinal Pigment Epithelium Injury in Pentosan Polysulfate Exposure: Morphologic Changes, Phagocytic Deficits, and Mitochondrial Dysfunction

**DOI:** 10.64898/2026.03.28.715018

**Authors:** Archeta Rajagopalan, Ganesh Satyanarayana, Rabina Kumpakha, Sakshi Shiromani, Jeffrey H. Boatright, Nieraj Jain, Sayantan Datta

## Abstract

Pentosan polysulfate (PPS) is a semisynthetic sulfated polysaccharide that was approved by the United States Food and Drug Administration (FDA) for treatment of interstitial cystitis (IC). A 2018 study by our group described a vision-threatening macular toxicity associated with long-term use of PPS. However, given the relatively recent characterization of PPS maculopathy, we have limited knowledge of its pathophysiology. The present study therefore investigated the pathophysiology of PPS maculopathy in a cell culture model, assessing impacts of PPS exposure on morphology and mitochondrial function. We treated ARPE-19 cells with increasing doses of PPS and investigated both mitoprotective and cytoprotective mechanisms, mitochondrial reactive oxygen species production (ROS) and respiration, cellular structure, and retinal pigment epithelium (RPE) dysfunction through phagocytosis assays. We found that PPS increased mitochondrial superoxide accumulation and that increased doses of PPS impaired basal and maximal respiration in a Seahorse assay without the expected response of increases in the cellular energy sensor pAMPK. PPS exposure disrupted mitochondrial and cell protective mechanisms against ROS accumulation as assessed through examination of mitochondrial biogenesis markers PGC-1α and SIRT1 and autophagy markers LC3 and p62. PINK1 expression increased with increasing duration of exposure to PPS. Further, we found that PPS led to functional and structural changes to RPE cells, which exhibited an increase in cell aspect ratio and impaired phagocytosis with higher doses of PPS. Lastly, we found an increase in cell death in response to higher doses of PPS, evident through ethidium homodimer cell viability assays. Taken together, our study shows PPS exposure has profound effects on RPE viability and function through impairment of mitochondrial respiration and mito- and cyto-protective mechanisms and highlights mitochondrial insult as a potential focus of future PPS research.

## 1. Introduction

Pentosan polysulfate (PPS) is a semisynthetic sulfated polysaccharide that was approved in 1994 by the United States Food and Drug Administration (FDA) for treatment of IC, also known as bladder pain syndrome.^1^ A 2018 study by our group described a vision-threatening macular toxicity associated with long-term use of PPS.^2,3^ Given widespread global use of the drug spanning several decades, many thousands of individuals are thought to be at risk.

Despite relatively preserved visual acuity, patients with PPS maculopathy typically present with prolonged dark adaptation and difficulty reading.^2^ Fundus examination may reveal subtle macular pigmentary changes, while fundus autofluorescence (FAF) and near-infrared imaging (NIR) better reveal the extent of disease.^4^ Studies have shown this disease to be a dynamic process, with macular pigmentary changes leading to atrophy and progressive vision loss at later stages.^4^ Given the relatively recent characterization of PPS maculopathy and our limited knowledge of its pathophysiology, there are no known treatments to halt the disease progression.

Clinical studies suggest that PPS may lead to a primary injury to the RPE.^5^ Given similar retinal findings to mitochondrial maculopathies such as maternally inherited diabetes and deafness syndrome, it is speculated that the cellular injury may be mediated by a primary insult to RPE mitochondria.^6,7^

Mice treated with PPS show reduced RPE function and aberrant morphology.^8^ Other studies have used cell culture models to assess the effects of PPS treatment on RPE proliferation and migration.^9,10^ Taken together, these studies show PPS has significant effects on structure and function of the RPE.

The present study investigated the pathophysiology of PPS maculopathy in a cell culture model, assessing impacts on morphology and mitochondrial function. Based on previously published data, we treated the cells with PPS for 72 hours, and chose a range of PPS doses to investigate dose-dependent effects.^9,10^ We investigated both mitoprotective and cytoprotective mechanisms, mitochondrial reactive oxygen species production (ROS) and respiration, cellular structure, and RPE dysfunction through phagocytosis assays in response to increasing PPS doses.

## 2. Results

### 2.1 PPS leads to mitochondrial ROS accumulation and disruption of mitochondrial respiration

Given the well-documented role of mitochondrial dysfunction in the RPE in retinal degeneration, we initially looked at the effects of PPS on mitochondria.^11,12^ Using live cell imaging and MitoSox staining, we found a significant increase in mitochondrial superoxide accumulation at PPS 1.0 mg/mL (p<0.05) and 10 mg/mL (p<0.0001) doses (Figure 1A and 1B). Based on this finding, we conducted a Seahorse assay to assess changes in mitochondrial respiration in PPS-treated cells. This assay revealed that PPS exposure leads to decreased maximal respiration across time points for all doses of PPS (Figure 2A). Upon further analysis at 72-hour treatment with PPS, we found significantly decreased maximum ATP production (p<0.01) and decreased maximal respiration (p<0.01) for all concentrations of PPS, and a decrease in basal respiration at 10 mg/mL PPS treatment (p<0.0001), Figure 2B).

**Figure 1.**
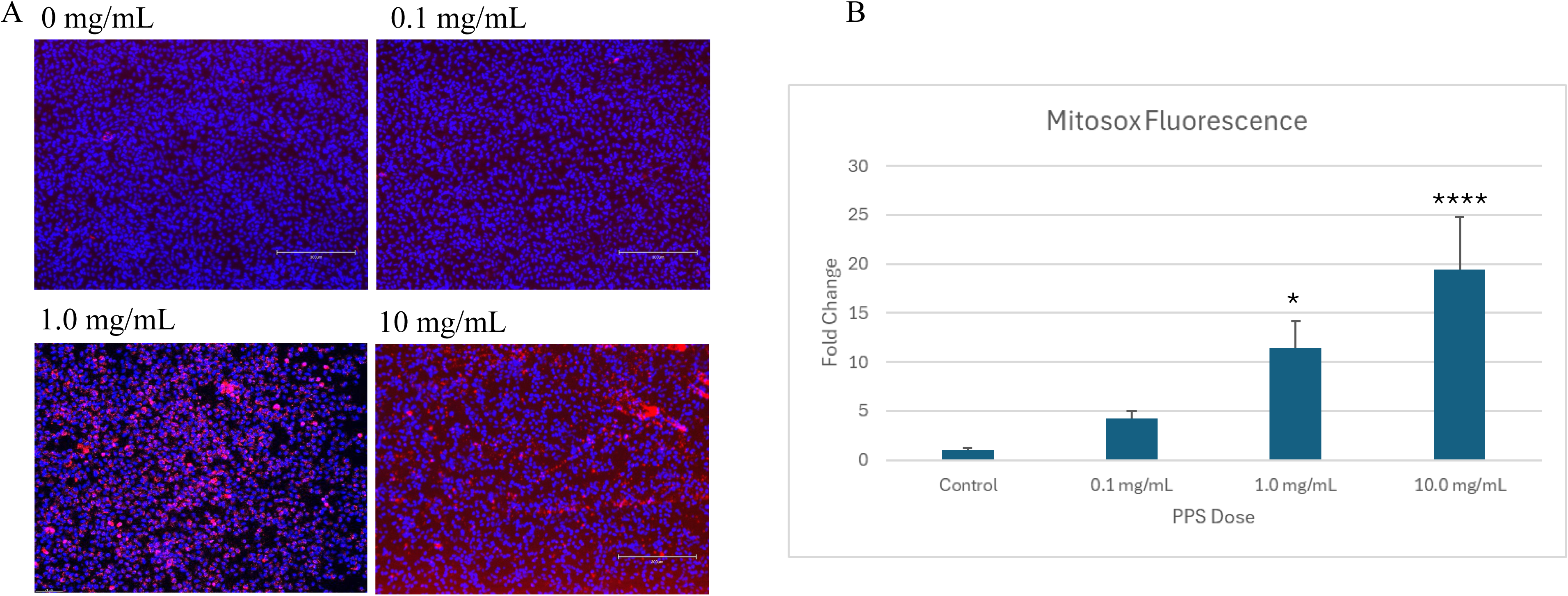
Mitosox (red) and Hoescht (blue) stain in ARPE-19 cells treated for 72 hours with increasing concentrations of PPS imaged with Leica Mica microscope at 10X magnification (A). Cells were imaged in live cell imaging buffer, and fluorescence was detected at excitation/emission of 396/610 nm. Red fluorescence was quantified using ImageJ and normalized to Hoescht stain. Fold change compared to the untreated control was calculated at each concentration for 3 biological replicates per concentration. Significance is reported with respect to control. Data shows significant increase in Mitosox at PPS 1.0 and 10.0 mg/mL concentrations (B).

**Figure 2.**
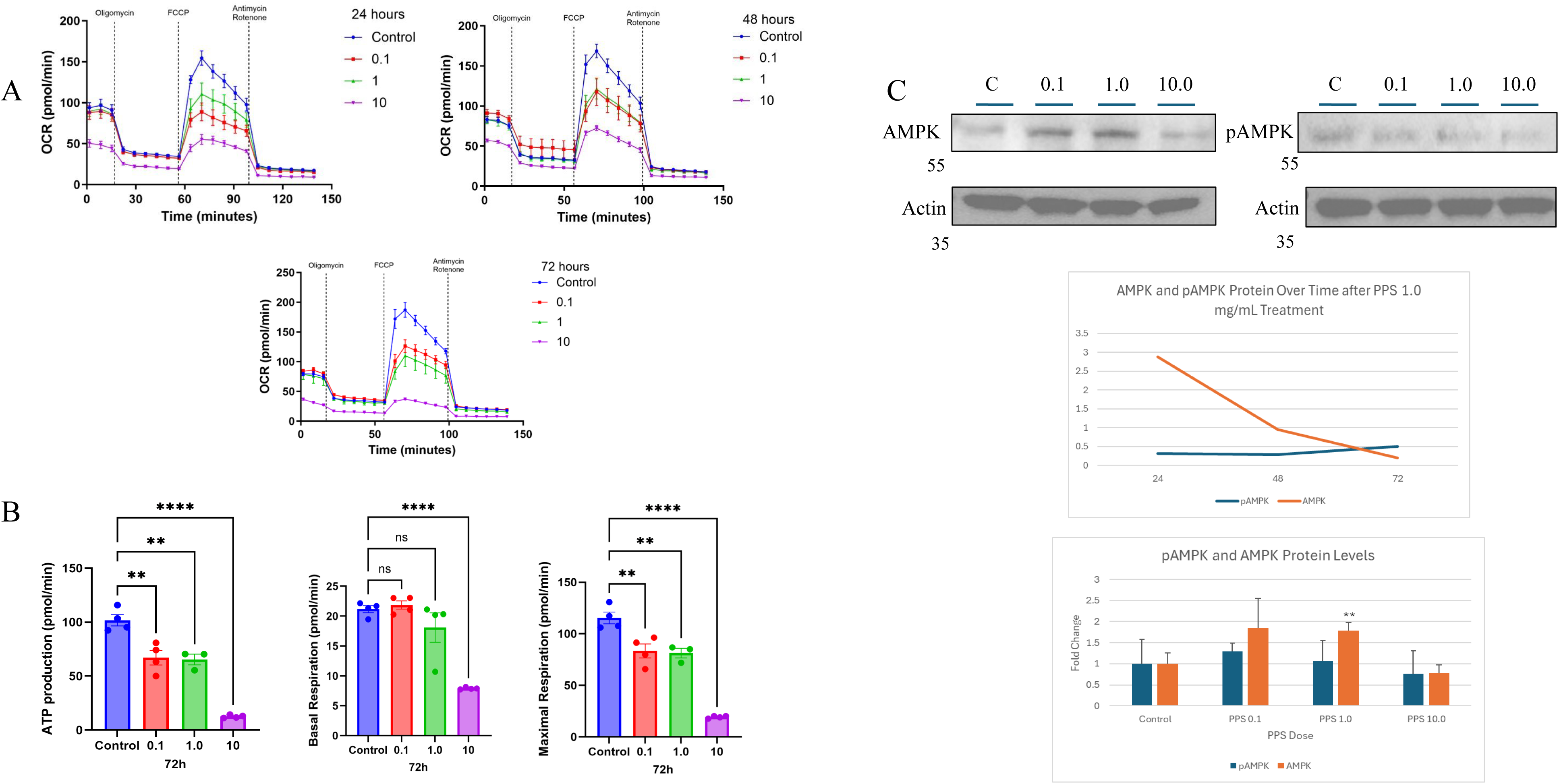
Seahorse experiment conducted for ARPE-19 cells treated with PPS 0, 0.1, 1.0, and 10 mg/mL at 24, 48, and 72 hours. Graphs show large decrease in respiration at PPS 10 mg/mL across time points. At 24 hours, PPS 0.1 shows larger insult to respiration than PPS 1.0 mg/mL, however the doses perform similarly at 48 hours, and the trend flips at 72 hours (A). ATP production, basal respiration, and maximal respiration obtained from Seahorse experiment with PPS 0, 0.1, 1.0, and 10 mg/mL at 72 hours showing significant decrease in ATP production at all concentrations of PPS, significant decrease in basal respiration at 10 mg/mL PPS, and significant decrease in maximal respiration at all concentrations of PPS (B). Western blot analysis of pAMPK and AMPK with corresponding **a**ctin with increasing concentrations of 72-hour PPS-treated ARPE-19 cells. Quantification with ImageJ and normalization to actin reveals significant increase in AMPK expression at PPS 1.0 mg/mL, without any increases in phosphorylated AMPK. Analysis of the time course of AMPK and pAMPK expression shows decrease in AMPK with longer treatment of PPS 1.0 mg/mL (C).

Western blot analysis followed with densitometry of cellular energy sensor AMPK and pAMPK (normalized to actin immunosignal) showed a trend of increased total AMPK at PPS 0.1 mg/mL and a significant increase at PPS 1.0 mg/mL (p<0.01), with no change at 10 mg/mL PPS treatment. There was no observed change in the active form, pAMPK, at any concentration.

These findings mirror the trend seen in the Seahorse data across concentrations in which the capacity for respiration in PPS-treated cells is diminished (Figure 2C).

### 2.2 PPS exposure leads to disruption of mitochondrial and cell protective mechanisms against ROS accumulation

We further explored the impact of PPS on mitochondrial function by observing the impacts of PPS on cell and mitochondrial protective mechanisms, namely mitophagy, mitochondrial biogenesis, and autophagy. qPCR analysis revealed a significant decrease in PGC-1α, a gene involved in the process of mitochondrial biogenesis,^13^ at a dose of 0.1 mg/mL PPS (p<0.01) and a significant increase at a dose of 10.0 mg/mL PPS (p<0.01). We also observed a significant increase in SIRT1, a gene involved in mitochondrial biogenesis and an activator of PGC-1α,^13^ expression at a dose of 1.0 mg/mL PPS (p<0.05) and at 10.0 mg/mL PPS (p<0.01). We did not observe any changes in PINK1, a mitochondrial serine/threonine-protein kinase that initiates mitophagy,^13^ between concentrations of PPS at 72 hours of treatment (Figure 3A). However, when observing changes in PINK1 expression at different durations of treatment with PPS, there was a significant increase in PINK1 expression between 24- and 72-hours treatment (p<0.001) and between 48 and 72 hours of treatment (p<0.0001) with PPS 1.0 mg/mL (Figure 3B). Western blot analysis of LC3 and p62, both involved in mitophagy,^14^ revealed increased expression at PPS 0.1 mg/mL and decreased expression at PPS 10 mg/mL (Figure 3C). Analysis of TOM20, an outer mitochondrial membrane protein used as a surrogate for total mitochondria in the cell,^15^ showed a parallel increase at 0.1 mg/mL PPS and a decrease at 1.0 mg/mL PPS (Figure 3C).

**Figure 3.**
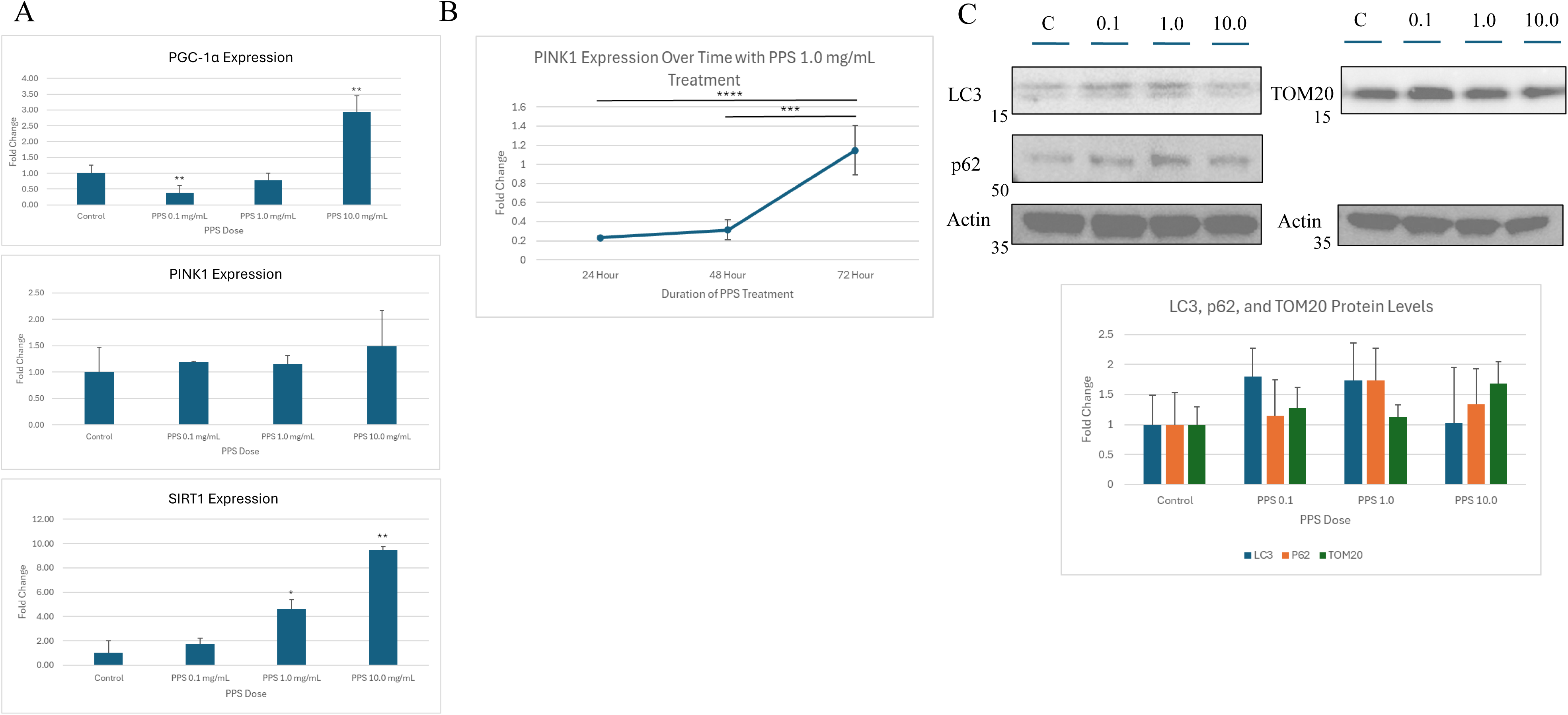
qPCR gene expression analysis conducted for ARPE-19 cells treated with PPS 0, 0.1, 1.0, and 10 mg/mL for 72 hours for PGC-1α, SIRT1, and PNK1. Significance is reported with respect to untreated control. Results show significant increase at PPS 10.0 mg/mL for PGC-1α and SIRT1 expression, and no significant change in PINK1 expression across concentrations (A). PINK1 expression at 24, 48, and 72-hour treatment time points with PPS 1.0 mg/mL relative to untreated control. There is significantly increased PNK1 expression with longer exposure duration (B). Western blot analysis conducted for LC3, p62, and TOM20 with corresponding actin showing trend of increasing LC3 and p62 at PPS 0.1 and 1.0 mg/mL with subsequent decreases at PPS 10.0 mg/mL (C).

### 2.3 PPS causes changes to RPE cell morphology with transition to EMT-like state

Based on the findings by Giradot et al., we explored changes in cell morphology in response to PPS dose.^8^ Brightfield microscopy showed a significant increase in cell aspect ratio at PPS 1.0 mg/mL (p<0.001) and PPS 10.0 mg/mL (p<0.0001) compared to untreated cells (Figure 4A). Further, qPCR analysis revealed a significant increase in ZEB1 (p<0.001) and TWIST (p<0.01), transcription factors that are key drivers of the epithelial-mesenchymal transition (EMT),^16,17^ at a dose of PPS 10.0 mg/mL (Figure 4B). Interestingly, coupled with this finding is a relatively stable level of vimentin, a cytoskeletal protein used as a biomarker for EMT,^16^ across doses on Western blot analysis (Figure 4C).

**Figure 4.**
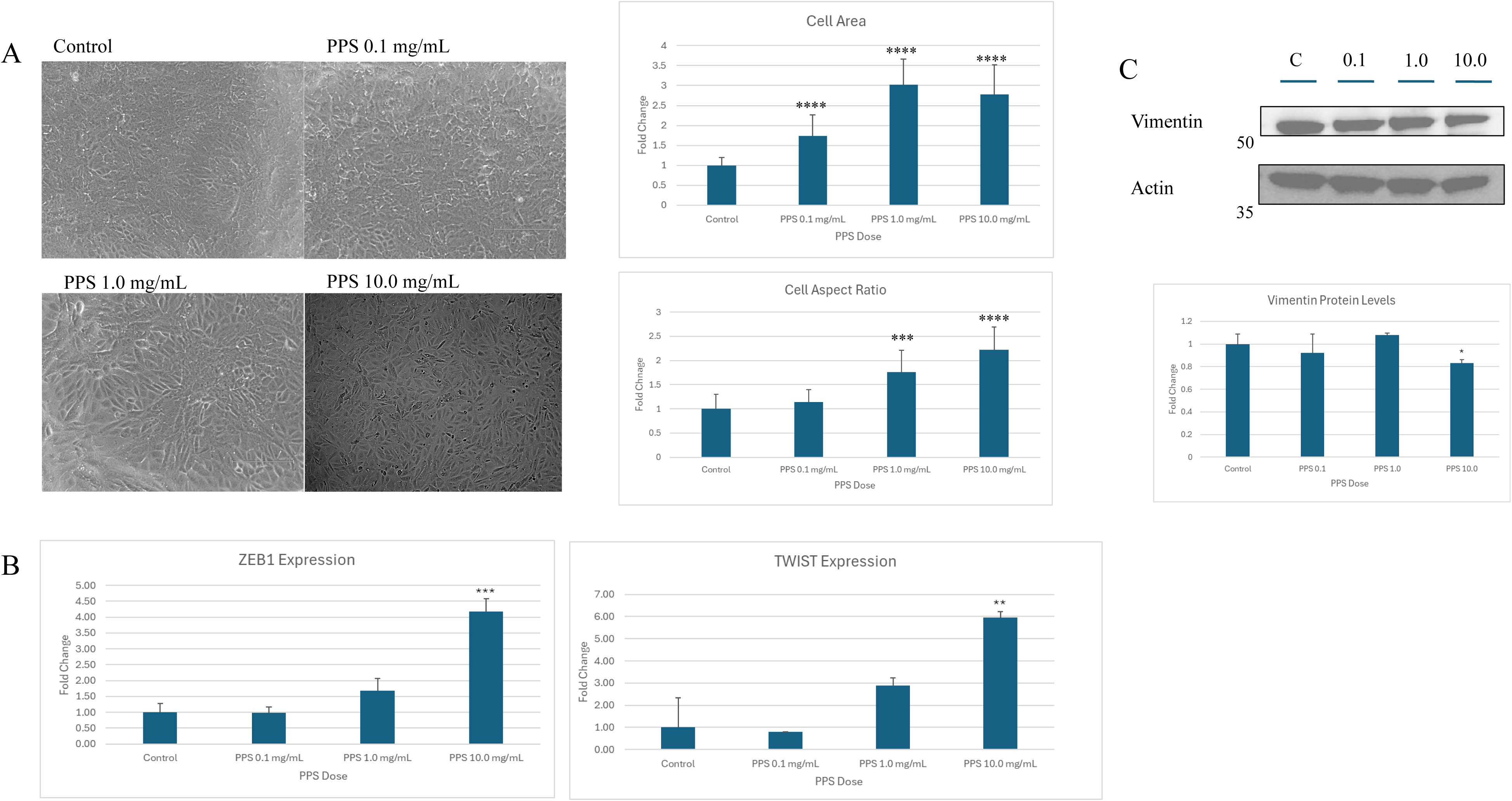
Brightfield imaging of ARPE-19 cells treated with PPS at 0, 0.1, 0.5, 1.0, and 10mg/mL at 72-hours of treatment. The cell outline was traced and parameters such as aspect ratio were quantified with ImageJ. Significance is reported with respect to control and shows a significant increase in cell aspect ratio at all doses of PPS (A). qPCR analysis of ZEB1 and TWIST expression in response to increasing PPS dose shows a significant increase in ZEB1 and TWIST expression at PPS 10.0 mg/mL. Significance is reported with respect to control (B). Western blot analysis of vimentin in response to increasing PPS dose showing a significant decrease with PPS 10 mg/mL dose (C).

### 2.4 PPS exposure causes functional changes in the RPE, leading to impaired phagocytosis capabilities

Previous published abstracts on the effect of PPS on ARPE-19 cells showed impaired phagocytosis capabilities of RPE after feeding with photoreceptor outer segments.^9^ We sought to replicate this finding using a pHrhodo assay. We find a decrease in phagocytosis of the pHrhodo particles at higher doses of PPS (Figure 5, p<0.0001), indicative of impaired phagocytosis in these cells.

**Figure 5.**
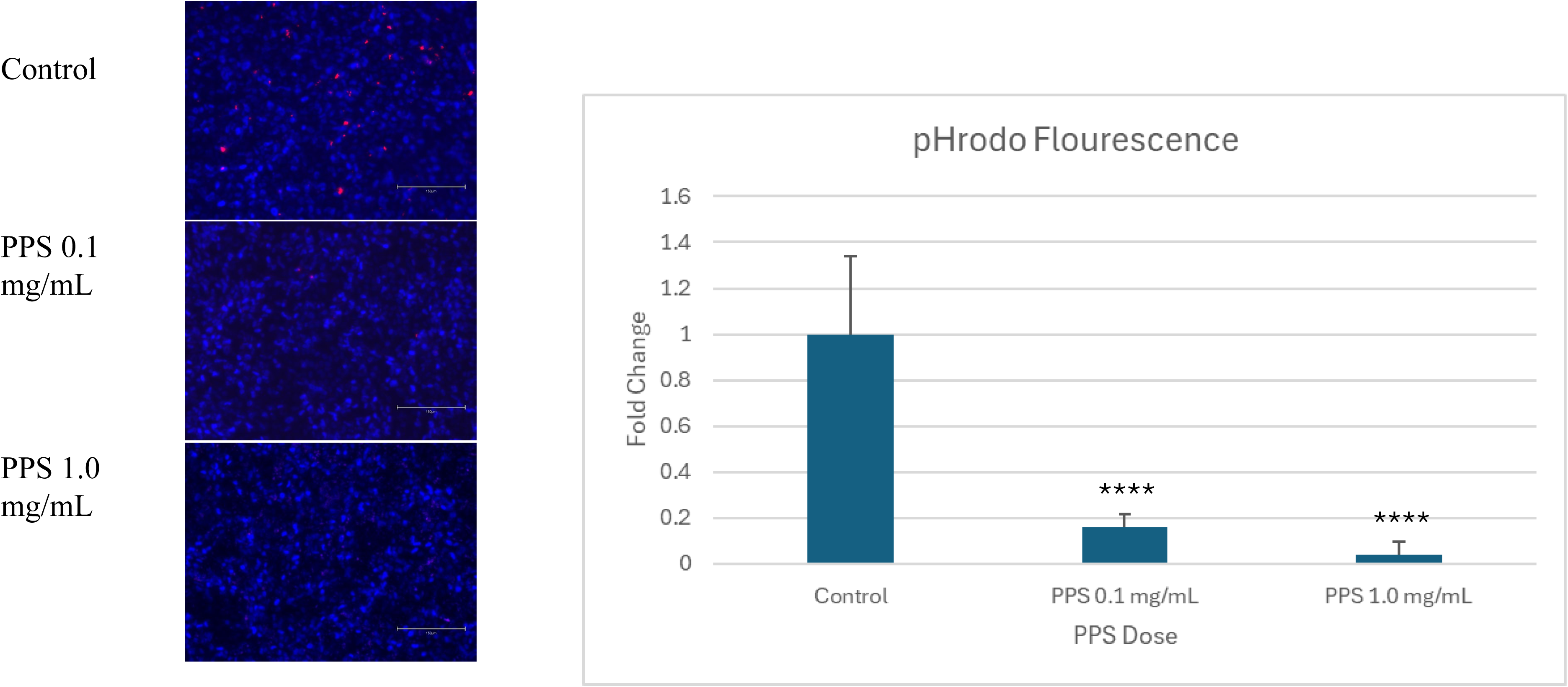
pHRhodo stain (red) and Hoescht (blue) in ARPE-19 cells treated with increasing doses of PPS. Cells were imaged in live cell imaging buffer, and fluorescence was detected at excitation/emission of 560/585 nm. Significance is reported with respect to control.

### 2.5 Exposure to higher PPS concentrations lead to cell death

Based on the accumulations of ROS in the mitochondria and evidence of impaired cellular protective mechanisms, we conducted an ethidium homodimer assay with live cells to assess cell death in response to PPS dose. Analysis showed an increase in ethidium homodimer accumulation at 1.0 mg/mL and 10 mg/mL doses of PPS, indicative of cellular death (Figure 6, p<0.0001).

**Figure 6.**
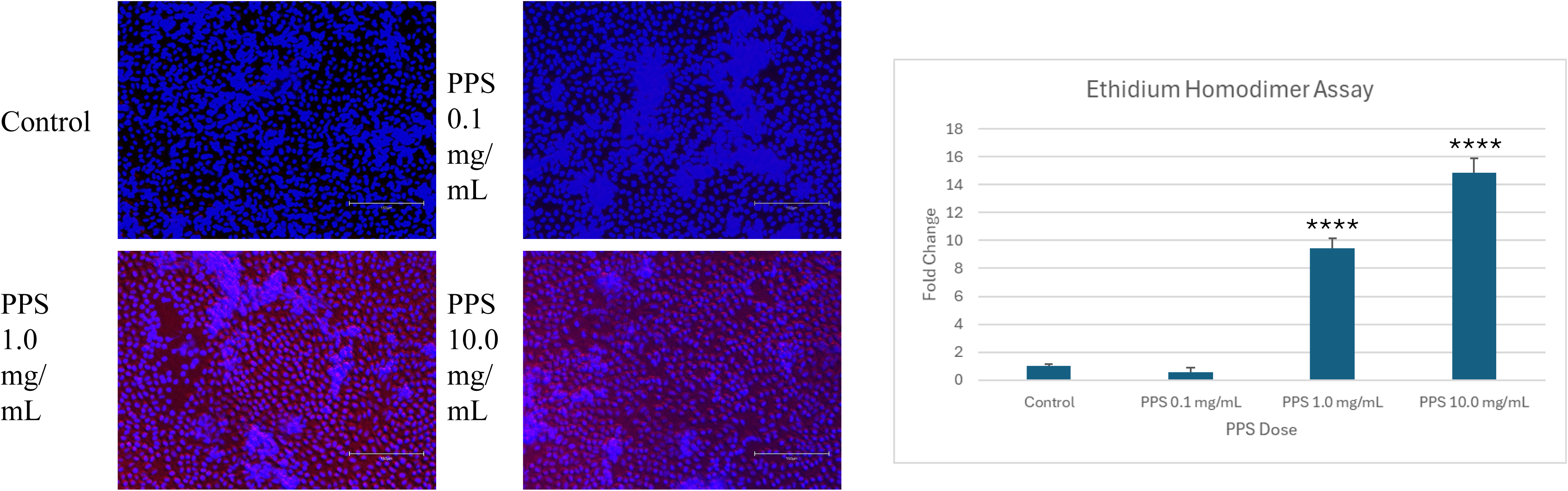
Ethdium homodimer (red) and Hoescht (blue) in ARPE-19 cells treated with increasing doses of PPS. Cells were imaged in live cell imaging buffer, and fluorescence was detected at excitation/emission of 528/617 nm. Significance is reported with respect to control.

## 3. Discussion

The present study explored mechanisms of toxicity in PPS maculopathy, highlighting a potential site of insult in RPE mitochondria.

Our results showed a significant impact at high doses of PPS on mitochondrial respiration and accumulation of mitochondrial superoxides. Paired with this finding, on an RNA level, we see cellular and mitochondrial protective mechanisms such as markers for autophagy and mitochondrial biogenesis at higher concentrations of PPS. Time course analysis also showed a significant increase in expression of mitophagy markers with longer duration of PPS treatment. Though the cell responded to PPS on a transcriptional level, we did not see the same trends on a translational level when observing protein markers for these processes. We also saw a large increase in cell death at higher concentrations of PPS, possibly due to these impaired mito- and cyto- protective processes. Further, we proposed that processes such as phagocytosis were impaired in our cell models at higher doses of PPS, possibly due to a lack of functional mitochondria for ATP production, as seen in our mitochondrial respiration assays. Finally, perhaps in response to this mitochondrial insult, we saw morphological changes in the cell, evident through the enlarged cell aspect ratio and elevated EMT markers at higher doses of PPS.

Previous studies describing the proposed mechanism of action of PPS maculopathy have shown similar findings. Recently, a study by Giradot et al. showed that mice treated with PPS had significantly reduced RPE function and aberrant cell morphology.^8^ In the study, mice were fed PPS-enriched chow at doses up to 2 g/kg/d, and serial electroretinography was performed.^8^ After 11 months of treatment, the mean scotopic a-, b-, and c-wave amplitudes of PPS-treated mice were significantly reduced compared with those of untreated mice.^8^ RPE flatmounts showed an increase in RPE cell area, solidity, and eccentricity in treated compared to untreated mice, mirroring our analysis showing increased cell aspect ratio in treated cells.^8^ Further, staining with alpha-catenin showed many areas of intracellular alpha-catenin staining across the RPE sheet in PPS-treated mice, with a significant increase in intracellular alpha-catenin staining in treated compared to untreated mice.^8^ Previous studies have shown this finding to be indicative of RPE stress, which is validated by the findings in our study.^18,19^

Other studies have used in vitro models to assess the effects of PPS treatment on cellular function. One study of human induced pluripotent stem cells and photoreceptor outer segments (POS) evaluated phagocytosis and digestion function in PPS-treated cells. The PPS treated RPE retained significantly more POS after 24 hours of phagocytosis than untreated controls, and the rate of POS digestion slowed at 24 and 48 hours of PPS exposure. These findings point towards impaired phagocytosis in PPS-treated cells, a finding further validated by our results.^9^ Another study showed PPS interferes with RPE proliferation and migration; after incubation for 72 hours with 1 mg/mL PPS, heparin binding EGF-induced proliferation and migration of ARPE-19 cells was significantly limited.^10^ Based on our evidence that PPS treatment disrupts mitochondrial function, it is possible this functional disruption ties to lack of ATP production. Taken together, these studies show PPS has significant effects on structure and function of the RPE at a molecular level.

In our exposure duration studies, we found that PINK1 expression increased with longer exposure to PPS. This finding mirrors that of studies that have observed PINK1 upregulation in response to oxidative stress conditions.^20^ Given PPS maculopathy is a progressive disease that is linked to long-term exposure to PPS, it is possible this increase in PINK1 expression is indicative of the toxicity to the mitochondria in response to long-term PPS exposure.

Our findings may explain clinical findings of phenotypic similarities between mitochondrial maculopathies and PPS maculopathy. Retinal changes present in patients with some mitochondrial diseases, including maternally inherited diabetes and deafness (MIDD), include perifoveal or peripapillary pigment changes on fundus examination, and speckled or reticular hyper-autofluorescence and hypo-autofluorescence along with areas of RPE atrophy on FAF.^6,7,21^ In this study, we chose PPS at concentrations of 0.1, 1.0, and 10 mg/mL and reported most of our findings with 72 hours of treatment. Our initial testing involved treating and testing cells at 24 and 48 hours of treatment, however we chose 72-hours as our primary time point as there were minimal differences in cell morphology and overall findings, and previous studies focused their work on this duration of treatment.^9,10^ Further, given PPS maculopathy is typically characterized by long-term exposure to PPS, we felt that a longer incubation period with the drug would better represent the clinical disease.

This study has several limitations. Firstly, we conducted this study in an ARPE-19 cell line. Future research should include studies in induced pluripotent stem cell-derived RPE cell lines to more closely resemble pathways of mature RPE cells *in vivo*. Further, while we tested various doses of PPS and its effect on the ARPE-19 cells, it is unclear what doses *in vitro* mirror those taken by patients, given differences in the half-life and metabolism of PPS in each setting. Similarly, patients with PPS maculopathy take PPS over the course of many years, often with a large cumulative exposure.^5^ While we attempted to mimic this long-term exposure with multi-day treatment, it is unclear whether this mimics long-term exposure *in vivo*.

The present study shows that PPS induces mitochondrial dysfunction in RPE cells through disrupting cellular energetics and overwhelming mitoprotective mechanisms, suggesting a possible way in which long-term exposure to PPS may be damaging the RPE.

## 4. Methods

### 4.1 Cell Culture

ARPE-19 cells were cultured as previously described.^22^ PPS was filtered and prepared at a 10 micromolar stock. Unless otherwise noted, for the following experiments, cells were treated with PPS 0, 0.1, 1.0, and 10.0 mg/mL concentrations for 72 hours and plated in 12-well plates.

### 4.2 Mitochondrial Assays

Cells were plated and treated in 96-well plates. Mitochondrial superoxide was assessed with MitoSOX Red (Thermo Fisher Scientific, M36008).

### 4.3 RNA extraction

Total RNA was extracted using the BioRad RNA Isolation kit. RNA quality was confirmed with the BioTek Synergy H1 Microplate Reader (Agilent Technologies).

### 4.4 Quantitative real-time RT-PCR (RT-qPCR)

Reverse transcription was performed using High-Capacity RNA-to-cDNA Kit (Thermo Fisher Scientific). RT-qPCR analyses were performed using primers listed in Table 1 with a Fast Universal PCR Taqman system (Thermo Fisher Scientific).

**Table 1.** Human primers used for qPCR analysis.

| Human Primers | Gene Taqman Probes |
| --- | --- |
| PINK1 | Hs00260868_m1 |
| GAPDH | Hs02786624_g1 |
| ZEB1 | Hs01566408_m1 |
| PPARGC1a | HS01016719_m1 |
| TWIST | Hs00361186_m1 |
| SIRT1 | Hs01009006_m1 |

### 4.5 Western blot analysis

Cell lysates were prepared in RIPA lysis buffer with protease and phosphatase inhibitor cocktail. Protein concentration was measured using DC Assay (BioRad, 5000111) and 5 ug samples were prepared using NuPAGE™ LDS Sample Buffer (4X) (Invitrogen). Samples were separated on 10% bis-tris SDS-PAGE gel and transferred to a nitrocellulose membrane. Membranes were blocked for one hour with 5% bovine serum albumin (BSA). Membranes were incubated with primary antibody (Table 2) at a concentration of 1:1000 overnight at 4°C, and the appropriate horseradish peroxidase conjugated secondary antibody was used at a concentration of 1:2000. Membranes were washed with 1x TBS and 0.1% Tween-20. Signals from western blots were detected with a chemiluminescence detection system (Pierce ECL or SuperSignal West Femto, Thermo Fisher Scientific). Blots were imaged with a ChemiDoc MP Imaging System scanner (BioRad). Band intensity is reported as arbitrary densitometric units.

**Table 2.** Antibodies used for Western blot analysis.

| Antibody | Concentration | Catalog No. | Vendor | Location |
| --- | --- | --- | --- | --- |
| Anti-MAP1/LC3B | 1:1000 | L7543 | Sigma-Aldrich | St. Louis, MO |
| Anti-VIM | 1:1000 | sc373,717 | Santa Cruz<br>Biotechnology | Dallas, TX |
| Anti-phospho-AMPK | 1:1000 | 2535S | Cell Signaling<br>Technology | Danvers, MA |
| Anti-AMPK | 1:1000 | 5831S | Cell Signaling<br>Technology | Danvers, MA |
| Anti-TOM20 | 1:1000 | sc-11415 | Santa Cruz<br>Biotechnology | Dallas, TX |
| Anti-E-cadherin | 1:1000 | 24E10 | Cell Signaling<br>Technology | Danvers, MA |
| Anti-p62 | 1:1000 | M162-3 | MBL Life Sciences |  |
| Goat anti-rabbit (IgG)<br>HRP | 1:2000 | Ab6721 | Abcam | Waltham, MA |
| Rabbit anti-mouse<br>(IgG) HRP | 1:2000 | Ab6728 | Abcam | Waltham, MA |

### 4.6 Seahorse Assay

Mitochondrial oxygen consumption rate from cell culture media was quantified with the Seahorse Bioscience XFe96 analyzer (Agilent Inc., Santa Clara, CA, USA) and the XF Cell Mito Stress Test. For Seahorse experiments, cellular respiration was analyzed by directly measuring the mitochondrial oxygen consumption rate (OCR) from the cell culture media using a Seahorse Bioscience XFe96 analyzer in combination with the Seahorse Bioscience XF Cell Mito Stress Test assay kit (103,015–100) as per the manufacturer’s protocol. Briefly, cells were plated at 40,000 cells/well and cultured on Seahorse XFe96 plates for 3 days. 24, 48, and 72 hours prior to analysis, cells were treated with 0, 0.1, 1.0, and 10.0 mg/mL PPS. On the day of analysis, cell medium was removed then changed to XF base medium supplemented with 10 mmol/L glucose, 1 mmol/L sodium pyruvate, and 2 mmol/ L glutamine (pH 7.4) followed by incubation at 37°C in a non-CO2 incubator for 1 h. The analyzer was then calibrated, and the experiment was run.

### 4.7 pHrhodo Assay

pHrodo assay was conducted per recommended protocol using pHrodo™ Red E. coli BioParticles™ Conjugate (Thermo Fisher Scientific, P35361). In short, 72-hour PPS-treated cells were treated with pHrodo particles in a 1:10 dilution in cell culture media and incubated at 37°C for 2 hours. Cells were washed 3 times with phosphate buffered saline (PBS) to remove excess undigested particles, and cells were imaged in live cell imaging buffer.

### 4.8 Assessing RPE Structure

Cells were plated in 8-well chamber slides and treated as above. Images were taken using EVOS M5000 Imaging System (Thermo Fisher Scientific) using brightfield at 20x magnification. The images were then analyzed using Image-J. After scaling the image, cells were manually outlined and cellular parameters including aspect ratio were measured. Fifty cells were outlined per image, and three biological replicates were included per treatment group. The average cell aspect ratio for each treatment group was calculated and normalized to the control to determine the fold change in cell aspect ratio.

### 4.9 Ethidium Homodimer Assay

Cells were plated in 96-well plates at a seeding density of 0.1 x 10^6^. Cells were treated once they reached confluence (0.4 x 10^6^) for 72 hours. Cell death was assessed using Ethidium Homodimer (Invitrogen, E1169) at 2.5 micromolar, incubated at 37°C for 30 minutes.

### 4.10 Statistical Analysis

The difference between groups was compared by Student’s t-test using GraphPad Software (San Diego, CA) after verifying that the data were normally distributed using the Kolmogorov-Smirnov Test of Normality. All data are expressed as the mean ± SEM or SD. Experiments were conducted in triplicate and performed at least 3 times.

**Supplemental Figure 1.** Full representative blots for each antibody used in analysis with corresponding expected molecular weight.

## Notes

Supported by the The Abraham J. and Phyllis Katz Foundation (JB); VA RR&D C9246C, I01RX002806, and I21RX001924 (JB); National Institutes of Health (NIH) R01EY028859 (JB), NIH R01EY031042 (JB); Foundation Fighting Blindness CD-C-0918-0748-EEC (NJ); Research to Prevent Blindness Challenge Grant (JB, NJ, SD); NIH P30EY006360 (JB, NJ, SD); NIH R00EY029010 (SD); Emory Eye Center (SD); Sitaraman family (NJ); Eberle family (NJ); Dobbs Foundation (SD)

### Competing Interest Statement

The authors have declared no competing interest.

